# Connectomic Flow Across Cortical Sensory Systems in Juvenile Fibromyalgia

**DOI:** 10.64898/2026.11.14.766636

**Authors:** Laura Martín-Herrero, Maria Suñol, Elisenda Bueichekú, Saül Pascual-Diaz, Tracy V. Ting, Jonathan A. Dudley, Catherine J. Jackson, Susmita Kashikar-Zuck, Robert C. Coghill, Jorge Sepulcre, Marina López-Solà

## Abstract

Adult fibromyalgia (FM) is characterized by altered functional connectivity between sensory, salience, and default mode networks (DMN), yet its early emergence and hierarchical organization remain unclear in Juvenile Fibromyalgia (JFM). We used stepwise functional connectivity (SFC) to detect early changes in the hierarchical integration of sensory signals toward salience and heteromodal networks in adolescents with JFM. Additionally, we assessed whether these alterations overlap with FM-associated gene expression from the Allen Human Brain Atlas (AHBA). Forty-six adolescent females with JFM (16.44±1.11 years) and forty-three healthy females (16.09±1.06 years) completed clinical questionnaires and underwent resting-state fMRI. Whole-brain voxel-wise SFC mapped connectivity patterns from primary sensory regions (visual, somatosensory, auditory) toward the DMN across increasing link-step distances. Regression models examined associations between symptom severity and SFC measures. Spatial overlap between functional findings and FM-associated gene expression profiles was also evaluated. Both groups showed a similar pattern, progressing from primary to secondary sensory regions, then to multimodal integration and salience areas, and finally to heteromodal DMN cortices. Greater symptom severity was associated with reduced second-step connectivity between secondary somatosensory cortices and the right anterior insula/operculum (T=5.11, pFWE<0.05). The neuronal transcription factor MYT1L showed relatively higher expression in AHBA in an overlapping region including the pars opercularis. Large-scale sensory-to-heteromodal functional architecture appears preserved in JFM. However, patients with greater symptom burden show reduced propagation of somatosensory information toward salience–interoceptive regions, suggesting altered multimodal processing. This disruption spatially overlaps with MYT1L expression, pointing towards a developmentally sensitive circuit vulnerability in severe patients.

## 1. Introduction

Chronic pain is conceptualized as a disorder of large-scale brain systems rather than reflecting altered nociceptive processing. Contemporary models emphasize hierarchical integration of sensory signals with salience, affective, and self-referential networks, which shape the subjective experience of bodily states and affective value assigned to sensory information [7,14,33,36]. In adult fibromyalgia, disruptions have been reported in interactions among somatosensory regions, salience circuitry, and heteromodal regions of the default mode network (DMN) [20,36]. Whether such alterations reflect early-emerging circuit vulnerabilities or adaptations to persistent pain remains unclear.

Adolescence represents a period of heightened neurodevelopmental plasticity, marked by ongoing refinement of long-range connectivity within fronto-insular and heteromodal networks. These systems support interoceptive awareness, salience detection (anterior insula), and the attribution of self-relevant affective value to moment-by-moment experience [6,22]. Investigating juvenile fibromyalgia (JFM), a chronic pain condition affecting 2–6% of youth, primarily adolescent girls, facilitates determining whether altered brain organization is present during this sensitive developmental window or emerges only after prolonged disease [19,45]. Notably, 80% of JFM patients continue to experience pain into adulthood, highlighting the importance of early mechanistic understanding [10,19].

Across adolescent and adult populations, fibromyalgia is associated with heightened unpleasantness to sensory stimuli and alterations in brain systems supporting sensory processing, salience attribution, and affective valuation [24,25,28,29]. Resting-state functional connectivity (rsFC) studies in adults have identified abnormal coupling between somatosensory regions, hubs of the salience network –the anterior insula, involved in sensory-interoceptive integration– and the DMN [7,20,33,36]. Additional work in both adult and juvenile samples has reported reduced connectivity within sensory cortices and altered brain responses to multisensory stimulation [15,24,25,28,29,36,41]. Despite these observations, it remains unclear whether fibromyalgia involves disruption across the topological transition from primary sensory regions to integrative salience-related and heteromodal networks, or whether dysfunction occurs at specific intramodal systems.

Large-scale brain organization is shaped by spatial gradients of gene expression that guide cortical specialization and network hierarchy [4,12,17,37,44]. Genetic studies have begun to uncover genes associated with fibromyalgia, including serotonin transporter gene (SLC6A4), transient receptor potential vanilloid channel 2 gene (TRPV2), myelin transcription factor 1 like gene (MYT1L), and neurexin 3 gene (NRXN3) [1,11,13]. MYT1L is of particular interest as a neuronal transcription factor that shapes neuronal identity and guides cortical circuit maturation [3,8,27], a process especially active during adolescence, a period of refinement of large-scale cortical circuits [22]. Examining normative transcriptomic maps, such as the Allen Human Brain Atlas (AHBA), could reveal whether the spatial expression of these genes aligns with cortical regions undergoing large-scale network maturation, a question that remains unexplored.

In this study, we applied stepwise functional connectivity (SFC), a graph theory approach mapping the topological reach and integration of connections across hierarchically organized brain systems, to test whether JFM is characterized by alterations in the transition from unimodal sensory processing toward salience-related and heteromodal integrative networks. We further conducted an exploratory analysis to investigate whether these connectivity alterations spatially overlap with genes previously implicated in fibromyalgia, using normative transcriptomic data as a proxy, thereby providing a framework for vulnerable circuits during adolescent chronic pain.

## 2. Methods

### 2.1. Participants

Eighty-nine female adolescents participated in the study, including 46 with juvenile fibromyalgia (JFM) (16.44 ± 1.11 years) and 43 healthy controls (16.09 ± 1.06 years) (demographic information appears in **Table 1**). This study is an extended analysis of a previously collected FIT-teens dataset [18], addressing a distinct question on stepwise functional connectivity and its relationship with transcriptomic organization. The study was preregistered on the Open Science Framework (https://doi.org/10.17605/OSF.IO/F2QUP). Ethical approval was obtained from the Institutional Review Board of Cincinnati Children’s Hospital Medical Center (2017-7771), and informed assent and parental consent were obtained. Further details are provided in the **Supplementary Methods 1**.

**Table 1.** Demographic and clinical characteristics of the study sample.

|  | **Healthy Participants (n=43)** | **JFM Patients (n= 46)** | **Statistics** |  |
|---|---|---|---|---|
| **DEMOGRAPHIC VARIABLES** | **Mean ± SD** | **Mean ± SD** | **T / X2** | **p-value** |
| Age (years) | 16.09 ± 1.06 | 16.44 ± 1.11 | 1.55 | 0.12 |
| Race (C / NC) | 2.06 ± 0.25 | 2.09 ± 0.42 | 0.28 | 0.77 |
| Yearly Household Income (1–7) | 5.16 ± 1.92 | 4.77 ± 1.95 | -0.93 | 0.35 |
| Education Level of the Primary Caregiver (1–5) | 4.09 ± 0.86 | 3.80 ± 0.84 | -0.97 | 0.33 |
| Education Level of the Secondary Caregiver (1–5) | 3.80 ± 0.85 | 3.79 ± 1.03 | -0.02 | 0.98 |
| **CORE JFM SYMPTOMS** | **Mean ± SD** | **Mean ± SD** | **T** | **p-value** |
| Fibromyalgia Symptom Severity | 1.16 ± 1.39 | 8.86 ± 2.13 | 19.01 | <0.0001 |
| Widespread Pain Index | 0.28 ± 1.08 | 11.35 ± 1.02 | 20.23 | <0.0001 |

Note: **Yearly household income** is shown using a scale of 1–7, where 1 = <\$24,999; 2 = \$25,000 to \$49,999; 3 = \$50,000to \$74,999; 4 = \$75,000 to \$99,999; 5 = \$100,000 to \$124,999; and 6 = \$125,000 to \$149,999; 7 > \$150,000. **Primary caregiver** refers to the person who is primarily responsible for meeting the child’s daily needs (mother in 74 cases, father in 6 cases); **Secondary caregiver** refers to the person who also plays a significant role in the child's care and well-being but may not have the primary responsibility (father in 67 cases, mother in 6 cases, not assigned for 7 cases). **Caregiver education level** is shown using a scale of 1–5, where 1 = less than high school; 2 = high school/GED;3 = partial college or trade school; 4 = college graduate; 5 = postgraduate degree. AASP, Adult Adolescent Sensory Profile; C, Caucasian; JFM, juvenile fibromyalgia; NC, Non-Caucasian; SD, standard deviation.

### 2.2. Clinical Measures

Developmentally appropriate and validated self-reported measures were used to assess core disease symptoms and functional disability. Participants completed the Pain and Symptom Assessment Tool (PSAT) [10], a standardized tool for the classification of FM in pediatric patients. This tool includes the Fibromyalgia Symptom Severity scale (FMSS), assessing symptoms experienced in the past week, and the Widespread Pain Index (WPI), which records body areas where participants experienced pain during the last three months. Both the FMSS and WPI are direct-scoring measures, where higher scores reflect either greater symptom burden and severity or pain in more body areas. These scoring properties have been validated in both adult and pediatric populations [10,45].

### 2.3. Fibromyalgia Associated Genes

*SLC6A4*, *TRPV2, NRXN3* and *MYT1L* are four potential genes involved in key pathophysiological processes that may contribute to the development and progression to chronicity of FM [11] **(Supplementary Methods 2.1)**. These genes were selected for spatial transcriptomic analysis using the AHBA to investigate whether their normative cortical expression profiles align with brain regions showing altered SFC in JFM. Further AHBA dataset details are provided in the **Supplementary Methods 2.2**.

### 2.4. MRI Acquisition and Preprocessing

We collected resting-state BOLD functional magnetic resonance imaging and structural T1-weighted data with a Philips Ingenia 3.0-Tesla MR System (Philips Healthcare) equipped with a 32-channel head coil at the Cincinnati Children’s Hospital Medical Center. Imaging data were preprocessed and denoised using standard pipelines from the CONN Toolbox-20.b running on MATLAB-R2021a (Math Works Inc). Imaging data acquisition and preprocessing are detailed in **Supplementary Methods 3**.

### 2.5. Network analysis

The focus of this study is to characterize altered sensory to multimodal interconnections in FM, hence, we applied advanced network connectomic tools to understand their transition across the cortical mantle. First, we created an association matrix that quantifies voxel-wise functional connectivity across the brain. Next, we derived the bipartite matrix to characterize connectivity specifically between two predefined sets of seeds regions, focusing our analysis in the brain areas that are more vulnerable in FM. Finally, we implemented SFC, a network approach that aims to characterize regions connected to specific seed brain areas at different levels of link-step distances [39].

#### Association Matrix calculation

Using a mask of 2585 voxels covering cortical gray matter, in line with our focus on cortical hierarchical organization, we computed voxel-wise association matrices for each participant by calculating the Pearson product-moment correlation coefficient (r values and their *p* values) for the time course of all brain voxels in a pairwise manner, obtaining individual-level r-value and *p*-value matrices. To perform this analysis, the pre-processed resting-state image of each participant was converted to an N-by-M matrix, where N represents the image voxels and M is the 559 acquisition time points. As a result, we obtained matrices of 2585 × 2585 (**Figure 1A**). We retained only positive correlations to ensure interpretability of connectivity weights and to reduce potential confounds related to preprocessing-induced anti-correlations and motion-related artifacts [31,32,38,43]. We corrected each individual’s connectivity matrix by controlling for the rate of false positives with a family-wise error rate (FWE) approach (pFWE corrected < 0.001 was applied to the *p*-values matrix, resulting in only r-values with corrected *p*-values).

**Figure 1.**
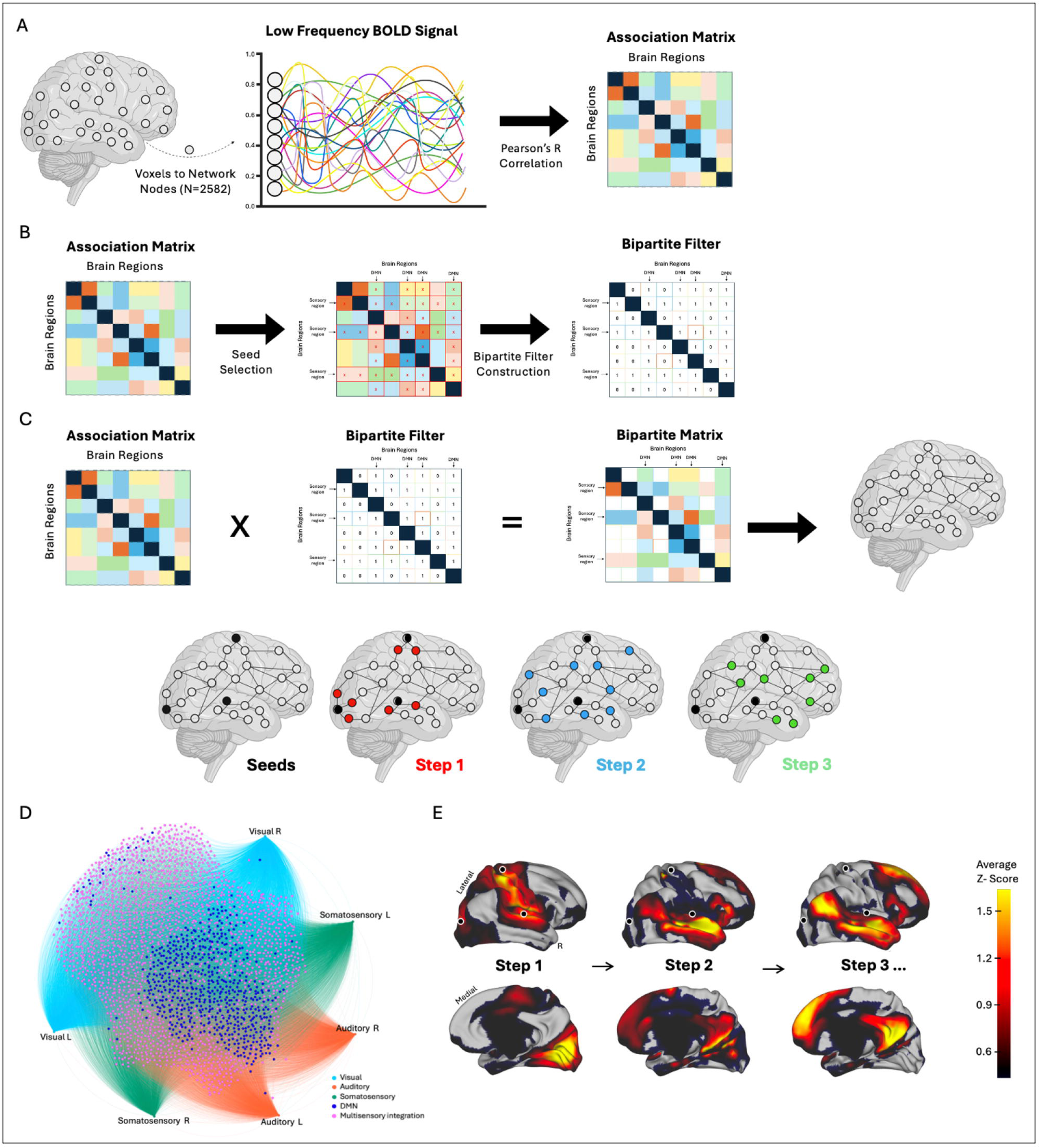
Pipeline of the stepwise connectivity analysis. **A)** An association matrix (A) was computed using Pearson correlation between voxels’ time courses within a cortical gray matter mask (number of voxels = 2582). **B)** The association matrix capturing whole-brain connectivity was used to select the values of the a priori seeds in primary sensory regions and the Default Mode Network (DMN). Based on these selections, a bipartite filter was constructed to map the connections between the two sets of regions, with rows representing the sensory regions and columns representing the DMN. **C)** The association matrix and the bipartite filter were combined using element-wise multiplication, resulting in the bipartite matrix that contains connectivity values for the selected seeds. The bipartite matrix is the starting point to obtain the cortical connectivity diagrams of *a priori* selected seeds using stepwise connectivity analysis. See Figure 2 for more details. **D)** Force-directed network map showing the characterization of stepwise connectivity analysis from *a priori* selected seeds to the DMN in JFM patients plotted using Gephi. In the force-directed network, each node (dot) represents a brain region from the bipartite matrix. The spatial layout is determined by a force-directed algorithm, which arranges nodes with stronger or more numerous connections closer together, thereby facilitating the visualization of symptom-related or functional clusters. **E)** Cortical maps showing the characterization of stepwise connectivity analysis from all *a priori* selected primary sensory seeds to the Default Mode Network in JFM patients plotted using workbench.

#### Bipartite matrix calculation

As FM has been previously related to alterations in the connectivity patterns of sensory systems and the DMN, we adopted a region-wise approach and created a bipartite matrix. We used two sets of seeds in this analysis: set 1 comprised six seeds in bilateral primary sensory cortices (visual, somatosensory, and auditory areas; extracted from Sepulcre et al. 2012)[39]; and set 2 corresponds to brain regions within the DMN (640 voxels, each voxel treated as a seed, from Yeo et al., 2011) [47]. To perform this analysis, we used a bipartite filter that encodes the adjacency relationship between two disjoint sets of vertices by retaining links between-sets (value of 1; e.g., visual to DMN) and removes connections within-set (value of 0; e.g., visual to visual) (**Figure 1B**). Finally, the bipartite matrix was obtained via element-wise matrix multiplication between the initial association matrix and the bipartite filter (**Figure 1C**), preserving its original size (2585 x 2585 voxels). The resulting matrix is a weighted connectivity matrix in which within-set connections are set to zero. For visualization purposes, the bipartite matrix was plotted using a force-directed network graph layout (Gephi software; ForceAtlas2 layout). A force-directed network map is a visualization that places connected nodes closer and unrelated nodes further apart, helping to show patterns and groups within complex networks (**Figure 1D**).

#### Stepwise Functional Connectivity analysis

As the focus of this study is to investigate changes in brain connectomics relevant to sensory-modal interconnections, we selected bilateral seed regions in the primary visual, primary auditory, and primary somatosensory cortices (Sepulcre et al, 2012). In SFC analysis, the degree of stepwise connectivity of a voxel *j* for a given step distance *l* and a seed area *i* 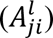 is computed from the count of all paths that (1) connect voxel *j* and any voxel in seed area *i*, and (2) have an exact length of *l*. Each SFC matrix *A_l_* of size m-by-m can be recursively represented as follows:

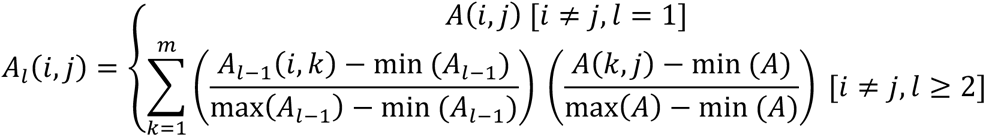

Here, *A_l_* is the weighted association functional connectivity matrix with a step distance of *l* after Fisher transformation. In the current analysis, we did a combined SFC analysis: we used the bipartite matrix as the input matrix for the SFC analysis and conducted the SFC analysis from all the seeds simultaneously. In other words, for each brain voxel, we calculated the number of links to all the six seed regions for a given step distance **(Figure 1C and Figure 1E)**. The SFC matrices were then normalized between 0 and 1, preserving the distribution of values intact while making them comparable across step distances. Then, we computed the SFC degree at each distance by summing the columns or rows of the corresponding SFC matrix. In this context, a high SFC degree represents a large number of paths connecting two voxels (the seed and target voxels) at a specific distance *l*. In **Figure 2** the SFC analysis is illustrated by the calculation of all link step distances between all the voxels of the brain and then extract the actual distance in a given length for a priori selected seed region within the network. It is important to note that while **Figure 2** uses an adjacency matrix (Adj) that is binary to illustrate the conceptual framework of path propagation in a simplified manner, the actual SFC analysis was implemented using the bipartite weighted matrix.

**Figure 2.**
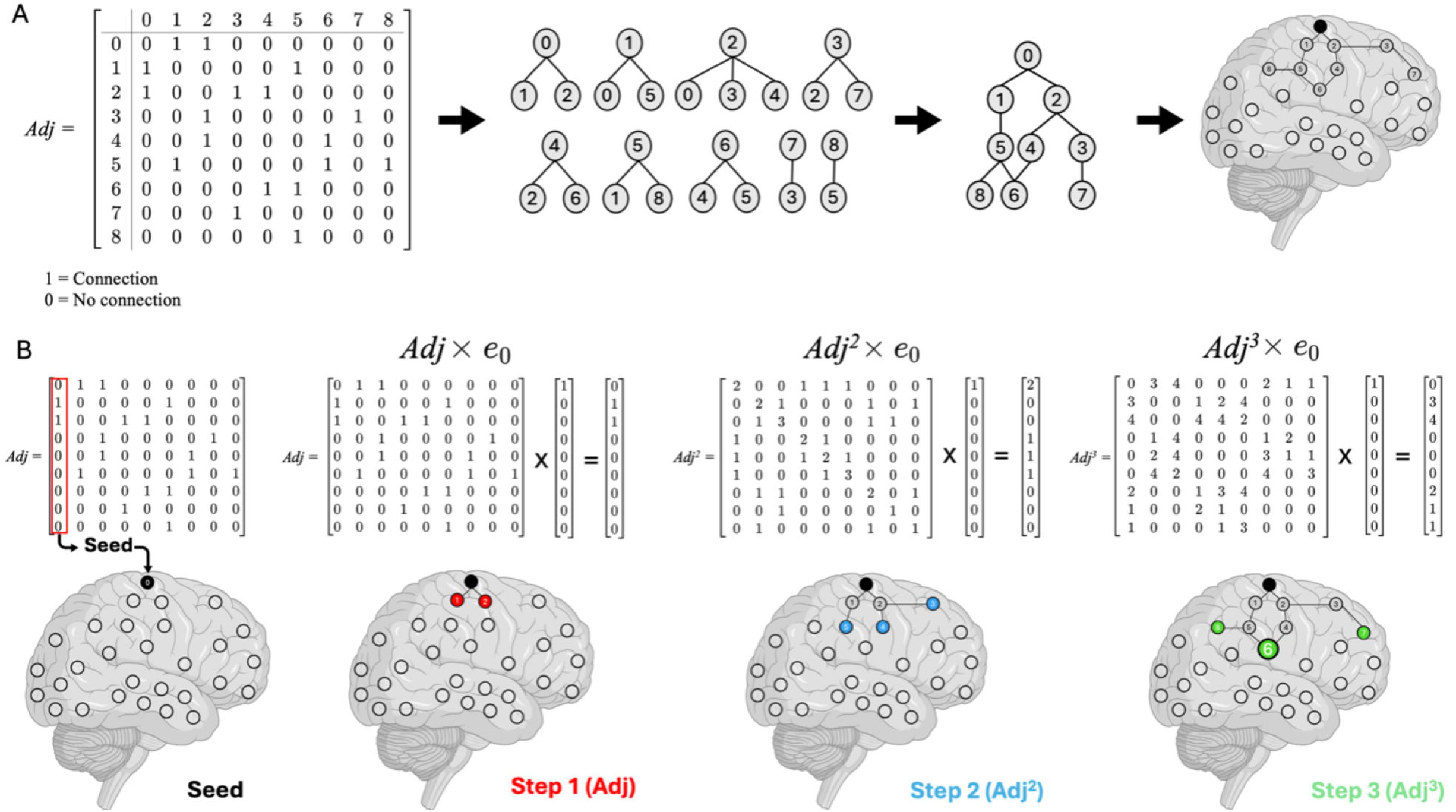
Stepwise connectivity analysis. **A)** Transformation from an adjacency matrix to a graph representation. From left to right: the adjacency matrix (Adj) encodes pairwise connections between brain regions, where a value of 1 indicates a connection and 0 indicates no connection; a graphical representation of all possible connections derived from adjacency matrix (Adj); a graph showing the overall network topology; and finally, the resulting connections mapped onto brain anatomy. This process illustrates how connectivity data can be progressively translated into interpretable network and anatomical representations. **B)** Illustration of stepwise connectivity from a seed region using weights of the adjacency matrix. The analysis starts from an initial vector (**e₀**), which is a binary vector indicating the seed (the region of interest) within the network. This vector is multiplied successively by the adjacency matrix (Adj) (i.e., Adj, Adj², Adj³), allowing the identification of regions that are connected to the seed through paths of increasing length (l), corresponding to 1-step (l=1), 2-step (l=2), and 3-step (l=3) connections. Each step identifies regions that are progressively further from the seed in terms of network topological distance. The corresponding brain maps visualize the spatial distribution of these connections, demonstrating the gradual spread of connectivity across the network. The size of node 6 is larger because it has a higher SFC degree at l=3; specifically, there are two distinct pathways of length 3 connecting the seed to node 6. In contrast, nodes 7 and 8 have only one pathway of the same length from the seed. The SFC degree reflects the number of unique paths of a given length that reach a node from the seed, highlighting its relative importance within the stepwise propagation of connectivity. **Note:** Figure 2 uses a simplified binary adjacency matrix for illustration. The actual SFC analysis uses the bipartite matrix derived from the association matrix, preserving connection strengths between sensory seeds and DMN. This weighted matrix guides the computation of stepwise connectivity and captures the true functional connectivity patterns.

We observed that beyond a link-step distance of three, the SFC pattern showed minimal changes in the spatial distribution, indicating topographic stabilization. Therefore, we only included maps from one to three steps in all our results. Using SPM12 (Wellcome Department of Imaging Neuroscience, London, England), we conducted one-sample t-tests to determine the main effects of connectivity within each group and link-step and a two-sample t-test was used to compare between-groups brain connectivity patterns (pFWE<0.05 at cluster-level and uncorrected p<0.001 at voxel-level).

After SFC analysis, we investigated the relationship between FM and brain connectivity. Using SPM12, we conducted a linear regression model between self-reported symptom severity scores (PSAT) and the SFC degree maps at each distance separately (pFWE<0.05 at cluster-level and uncorrected p<0.001 at voxel-level). The results were plotted using Connectome Workbench software (http://www.humanconnectome.org/software/get-connectome-workbench.html) **(Figure 1E)**.

### 2.6. Spatial associations between SFC and cortical transcriptomic information

Spatial associations between stepwise functional connectivity (SFC) maps and cortical gene expression were assessed using a validated neurogenetic pipeline (Sepulcre et al., 2018). Gene expression data from the AHBA were mapped to the Desikan–Killiany cortical atlas to examine spatial correspondence between fibromyalgia-related genes (SLC6A4, TRPV2, NRXN3, MYT1L) and regions showing altered SFC in juvenile fibromyalgia. Further details are provided in the **Supplementary Methods 2.3**.

## 3. Results

### 3.1. Sample characteristics and differences in FM severity

JFM patients and healthy controls were matched across all demographic variables, including age, race, annual household income, and the education levels of both primary and secondary caregivers. As expected, JFM patients reported significantly higher FMSS (t=19.01, p<0.001) and WPI (t=20.23, p<0.001) compared with healthy participants (**Table 1**).

### 3.2. Stepwise Connectivity Pattern in JFM Patients

We found no significant differences between JFM patients and healthy participants in SFC patterns **(Figure 3)**. In the first step, connectivity extended from primary sensory regions (visual, auditory and somatosensory) to secondary sensory regions within these modalities. In the second step, connectivity extended from the secondary sensory regions (visual, auditory and somatosensory) to multimodal (multisensory) integration brain areas. These include key regions such as the anterior insular cortex, the parietal operculum, the superior and the middle temporal gyrus, the posterior and the anterior cingulate cortex and the dorsolateral prefrontal cortex. In the third step, connectivity showed specific pathways from multimodal (multisensory) integration areas to heteromodal/association cortices, particularly the DMN.

**Figure 3.**
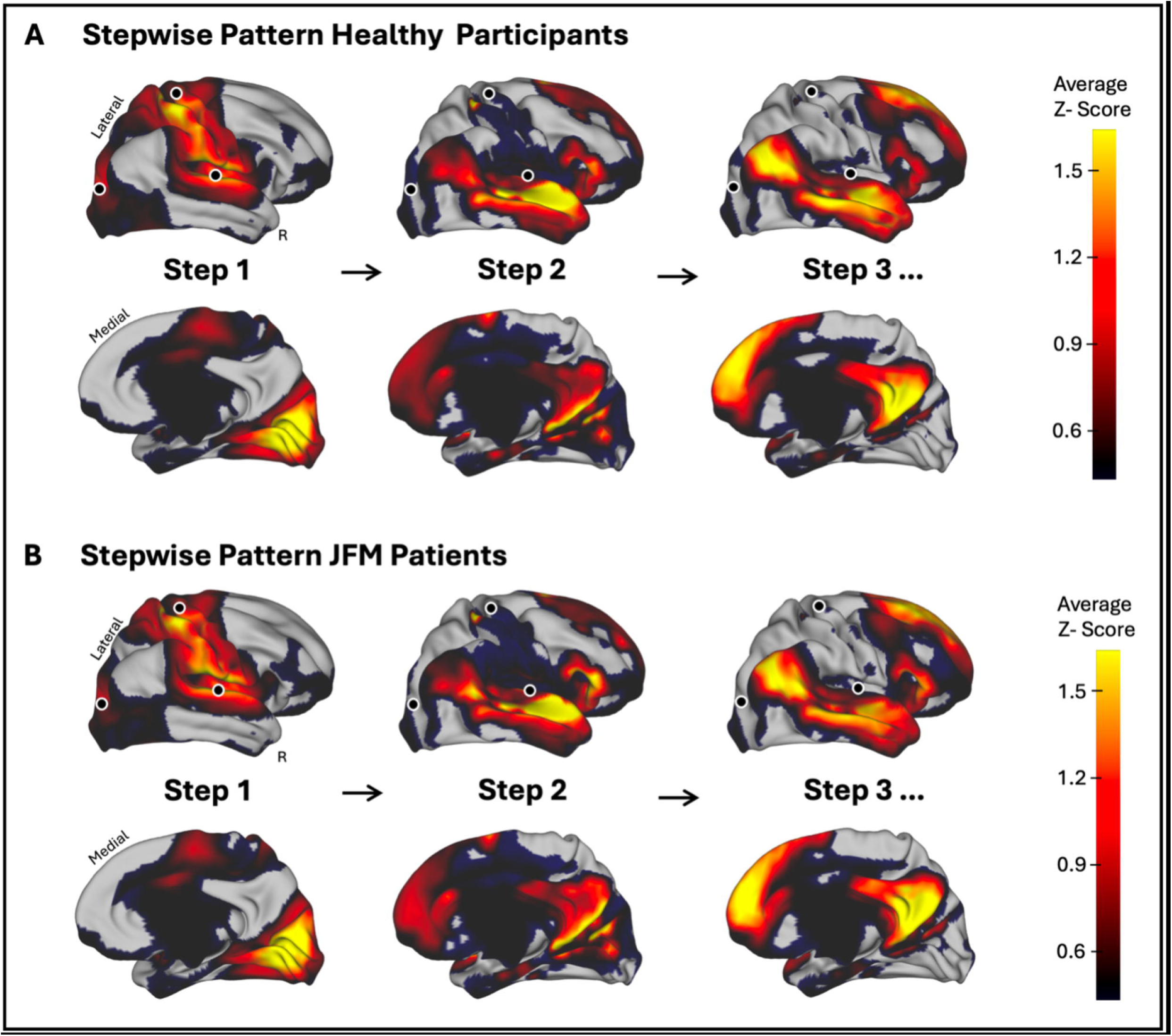
Resting-state connectivity patterns in healthy adolescents and adolescents with juvenile fibromyalgia. **A)** Mean cortical maps showing stepwise connectivity patterns in healthy participants. **B)** Mean cortical maps showing stepwise connectivity patterns in JFM patients. Values represent average Z-scores extracted from statistically thresholded connectivity maps (p < 0.001, FWE-corrected at voxel level in SPM12). Black regions indicate voxels that did not reach the statistical threshold and are shown for reference only. JFM, juvenile fibromyalgia; R, right.

### 3.3. Reduced Connectivity in JFM Patients with Higher Symptom Severity

JFM patients with higher symptom severity scores (measured using the FMSS) showed reduced second step connectivity between bilateral secondary somatosensory cortices to a cluster centered in the right anterior insula extending to the right pars opercularis (peak-level t=5.11, pFWE<0.05 at the cluster-level). Since our analysis was constrained by a bipartite matrix that models signal propagation specifically from the sensory regions to the DMN, this disruption suggests that the insula may serve as a critical information relay. Consequently, the reduced connectivity at this intermediary step may interfere with the flow of functional information toward the DMN, implying that greater symptom severity is linked to impaired integration along the somatosensory–insula–DMN trajectory **(Figure 4)**.

**Figure 4.**
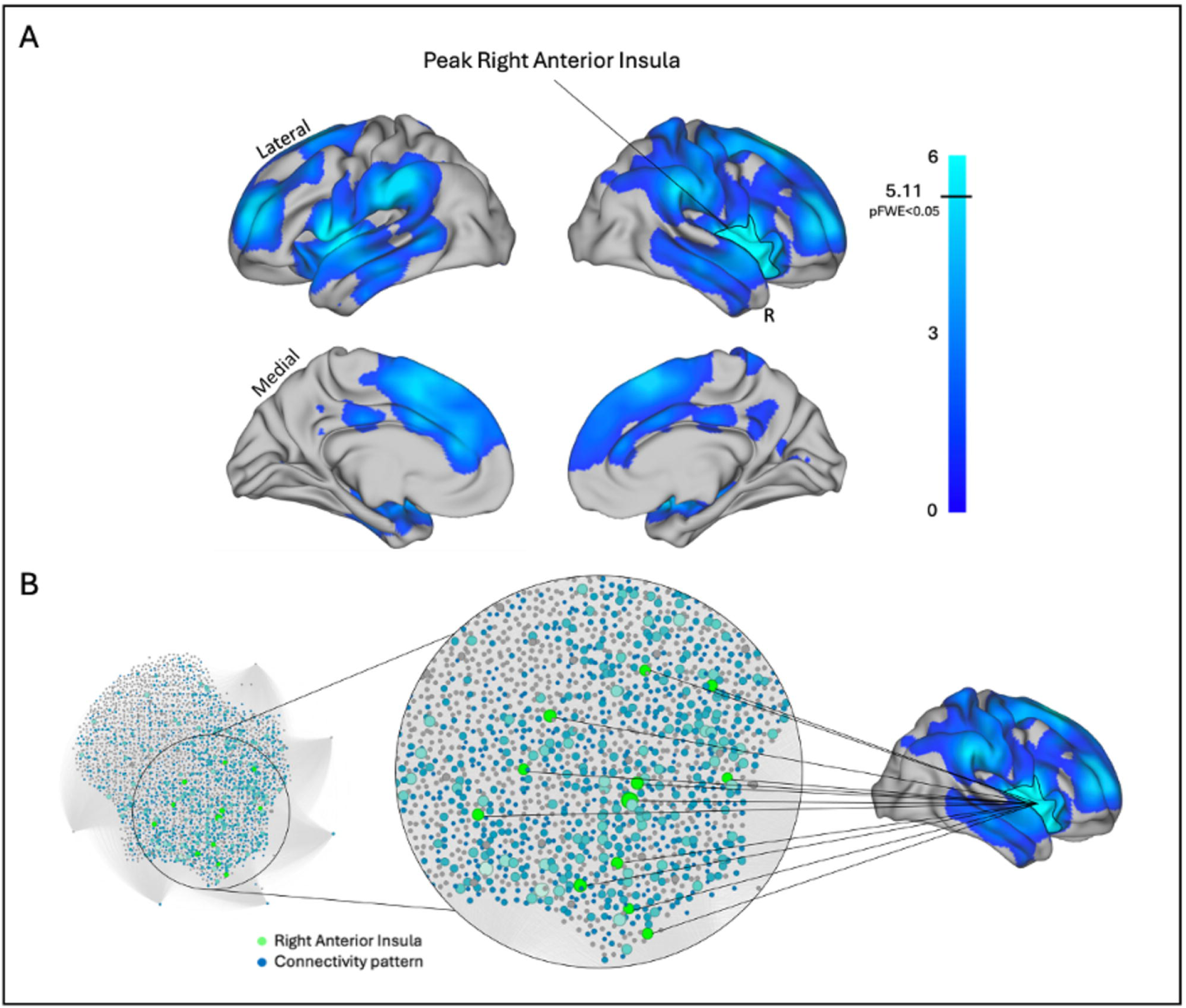
Higher symptom severity in juvenile fibromyalgia symptom is related to reduced connectivity in right anterior insula. Correlation map between stepwise functional connectivity results from the second step (connectivity extended from the secondary sensory regions to multimodal (multisensory) integration brain areas) and symptom severity in JFM patients**. A)** Cortical maps showing the weighted bipartite matrix of the second step correlated with higher symptom severity in JFM patients plotted using workbench. **B)** Combination of a force-directed network map and cortical surface maps, both illustrating the weighted bipartite matrix from the second step, which is correlated with higher symptom severity in JFM patients. The network map was generated using Gephi, while the cortical maps were plotted using Workbench. In the force-directed network, each node (dot) represents a brain region from the bipartite matrix. The spatial layout is determined by a force-directed algorithm, which arranges nodes with stronger or more numerous connections closer together, thereby facilitating the visualization of symptom-related or functional clusters. Node size reflects the SFC degree (i.e., the number of unique paths of a given length reaching the node from the seed), indicating the region’s overall involvement or influence within the network. Green nodes indicate the cluster that survived statistical correction (peak-level t = 5.11, pFWE < 0.05 at the cluster level), while blue nodes represent the remaining SFC connectivity pattern.

### 3.4. Link between Stepwise Connectivity and Cortical Gene Expression

The genetic expression of the FM-associated genes (*SLC6A4, TRPV2, MYT1L* and *NRXN3*) in each region of the Desikan–Killiany atlas is reported in the **Figure 5A**. *SLC6A4* is overexpressed in the right temporal pole (z = 4.99, temporal pole right upper bound z = 3.58), as its expression in that region exceeded the 1.96 SD threshold relative to the regional mean. Similarly, TRPV2 is overexpressed in the left lateral orbitofrontal cortex (z = 1.33, lateral orbitofrontal left upper bound z = 1.25), and *MYT1L* is overexpressed in the right pars opercularis (z = 2.05, pars opercularis right upper bound z = 2.03), based on the same within-region thresholding approach (**Figure 5B and Figure 5C)**.

**Figure 5.**
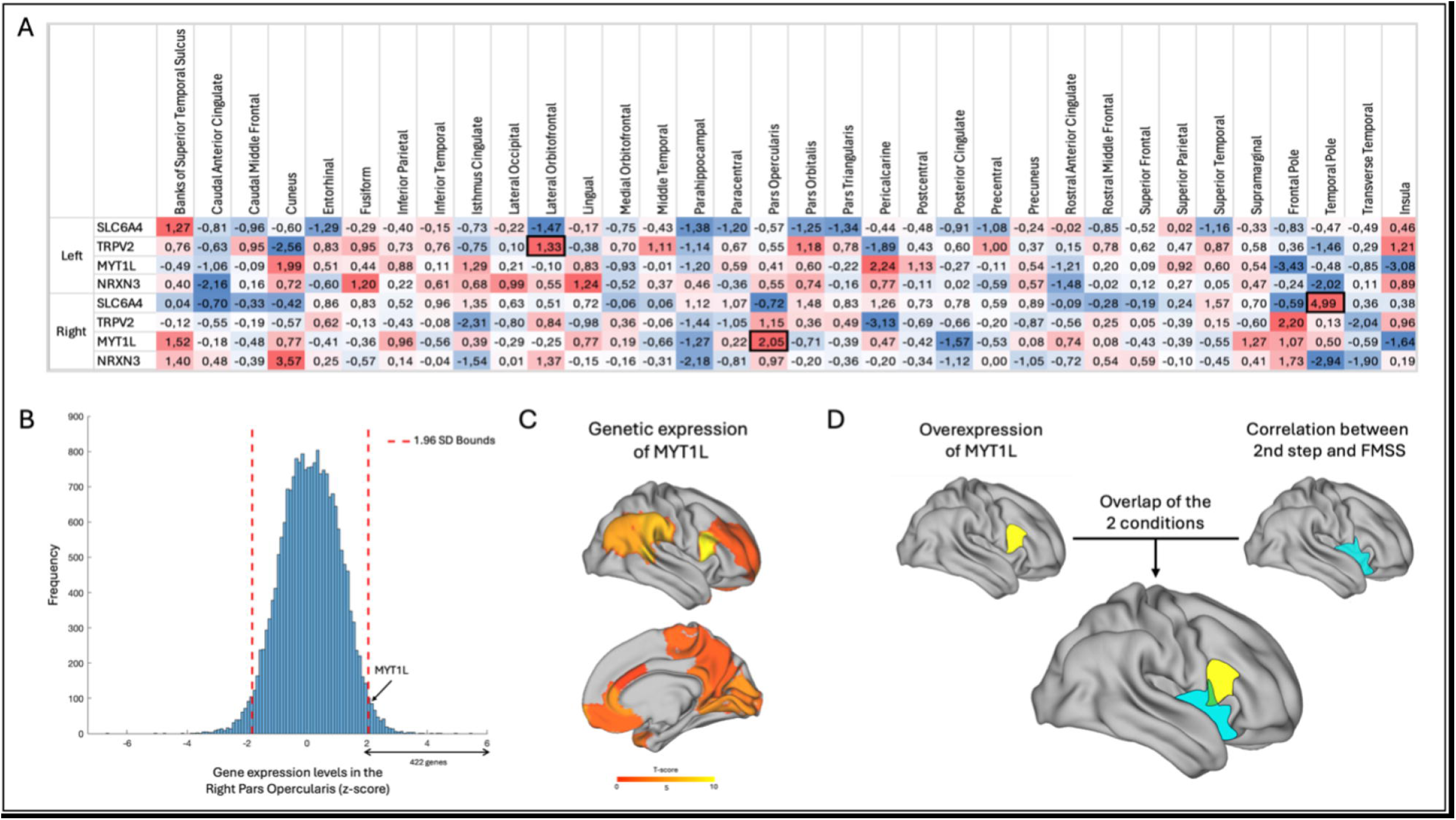
Spatial association of MYT1L expression with functional alterations in juvenile fibromyalgia. Association between gene expression and stepwise brain connectivity**: A)** Cortical gene expression of the FM associated genes (*SLC6A4, TRPV2, MYT1L* and *NRXN3*) across the human brain. Each value represents the normalized gene expression level (z-score) of a specific gene in each brain region, derived from the AHBA. Bolded values indicate significant overexpression, defined as z-scores exceeding +1.96, corresponding to the upper bound of the 95% confidence interval. **B)** Gene expression in the right pars opercularis region. **C)** Cortical map showing the spatial distribution of *MYT1L* gene expression. **D)** Cortical maps showing the spatial overlap between brain regions with overexpression of the *MYT1L* gene and the corrected SFC results reflecting associations between self-reported symptom severity scores and second step connectivity in patients. Note: Gene expression values are assessed relative to a region-specific null distribution rather than through across-region statistical testing; therefore, no correction for multiple comparisons across cortical regions was applied.

As an exploratory analysis, we assessed the spatial correspondence between the expression of FM-associated genes and the corrected SFC maps by directly spatially overlapping the two types of maps: gene expression maps and the maps linking self-reported symptom severity to second-step connectivity in patients. Specifically, patients with greater symptoms severity showed (i) reduced connectivity integration between somatosensory-right anterior insula/pars opercularis pathway, and (ii) overexpression of the *MYT1L* gene in an overlapping region of the right anterior insula/pars opercularis (**Figure 5D)**. This spatial overlay indicates a partial convergence between regions showing altered connectivity and regions of MYT1L overexpression. No spatial overlaps were found between the overexpression of *SLC6A4* or *TRPV2* and any of the other SFC maps.

## 4. Discussion

To our knowledge, this is the first study assessing multistep connectivity transitions (or connectivity flows) from sensory to association cortices in the brain of youth with chronic primary widespread pain. Adolescent females with JFM showed similar SFC patterns compared to healthy participants. Importantly, in the second step, where connectivity extended from the secondary sensory regions to multimodal (multisensory) integration brain areas, JFM patients with higher symptom severity showed less connectivity between bilateral secondary somatosensory cortices and the right anterior insula extending to the pars opercularis, suggesting that symptom severity in JFM is associated with disrupted information flow within the multimodal integration network. Furthermore, we observed an overexpression of the *MYT1L* gene that spatially overlapped with the right anterior insula/pars opercularis region. This suggests that the area of reduced connectivity integration in the somatosensory–insula pathway corresponds with an area of overexpression of *MYT1L* – in the right anterior insula/pars opercularis-, which may point to a possible convergence between gene expression and functional brain alterations in patients with FM.

The multimodal (multisensory) integration network processes the information from the different primary sensory cortices and connects them to the high-level brain networks involved in attention, emotional salience and executive control that operate in parallel at the top of the brain’s functional hierarchy [39]. This network gives rise to behavioral responses associated with the perception and interpretation of sensory stimuli. Interestingly, several studies report altered perception and processing of painful and non-painful sensory stimuli in chronic pain, and specifically in fibromyalgia -both adult and young patients-suggesting a potential alteration of the multimodal integration network already present in patients debuting at a young age and also when the disorder begins in adulthood [24,25,41,42]. Furthermore, abnormalities in multisensory integration may contribute to hypersensitivity across somatosensory, visual, auditory, and olfactory modalities, ultimately affecting pain perception and modulation [48]. These findings are consistent with the SFC pattern identified in both JFM patients and healthy participants of our study. Specifically, in the second step of the SFC analysis, we observed an extension of the connectivity from the secondary sensory regions (visual, auditory and somatosensory) towards the multimodal (multisensory) integration brain areas. Interestingly, in JFM patients with greater symptom severity we observed a hypoconnectivity between the bilateral secondary somatosensory cortices and a cluster in the right anterior insula extending to the pars opercularis. This suggests that such connectivity alterations do not characterize the patient group, but rather reflect individual differences related to symptom severity, consistent with previous findings from multisensory integration paradigms [29], and may serve as potential neural markers of clinical burden. Notably, the anterior insula has previously been identified as a central hub within the multimodal integration network, involved in the integration of perceptual inputs and cognitive control functions [9,39,40]. Moreover, previous studies have reported alterations in the right anterior insula in juvenile and adult FM [23,41]. This region has also been described as a switching core during pain processing that will relay sensory information into higher-order affective and cognitive modulation regions [30]. Altogether, these findings suggest that disrupted connectivity between the secondary somatosensory cortices and the anterior insula may impair effective multisensory integration and contribute to heightened pain sensitivity and symptom severity in adolescents with FM.

In further steps of the SFC pattern, connectivity extended across multimodal to heteromodal/association cortices, particularly the DMN. This is in line with previous studies in which SFC revealed patterns of sensory-specific connectivity showing a progressive organization from sensory to multimodal integration networks and ultimately involving DMN regions [35,39]. The DMN is associated with autobiographical, self, and social functions, and has also been link to pain modulation through descending inhibitory pathways [5,21]. In chronic pain conditions, rsFC studies have reported significant changes in DMN function [2,33]. Interestingly, our results showed that JFM patients with greater symptom severity exhibited reduced connectivity integration in the somatosensory-insula pathway, which is key step in the progression toward higher-order association regions, including the DMN. Although we did not observe direct alterations in the DMN, disruptions in earlier stages of the sensory-salience integration network may impair the engagement of heteromodal networks potentially contributing to the persistence and chronicity of pain in FM.

The implication for the potential cortical reorganization in FM, as observed in the present study and previous neuroimaging investigations [20,24,25,36,41,42], is also reported in genetics studies [13,34]. GWAS and Array Comparative Genomic Hybridization (aCGH) results point at a role for the CNS in FM genetic susceptibility [13]. Several investigations have shown changes in genes implicated in pain processing at the cortical level [16,26]. These findings provide converging evidence from both neuroimaging and genetics, suggesting that cortical dysfunction is a core component in FM pathophysiology. In this context, *MYT1L*, which has been previously linked to FM, may represent a gene of interest due to its involvement in various aspects of neurodevelopment, including neuronal identity, neuronal migration, maturation, and synapse formation [46]. We found that the overexpression of the *MYT1L* gene spatially overlaps with the hypoconnectivity observed between the secondary sensory areas and the right anterior insula/right pars opercularis in JFM patients with heightened FM symptom severity. This suggests a potential alignment between transcriptional architecture and functional connectivity alterations in FM. However, given the indirect nature of the datasets and the exploratory design of the analysis, these results should be interpreted cautiously. Overall, these observations may support a potential involvement of MYT1L-related processes in FM, although further validation using independent datasets and mechanistic approaches is required.

This study has several limitations. First, we exclusively enrolled cis-females. Thus, our findings cannot be generalized to other genders. This choice was based on our primary aim of investigating rsFC in a well-characterized sample that represents the population group predominantly affected by JFM. We also considered the potential influence of sex/gender differences in pain processing and brain developmental stage. Future studies should include adequately powered samples of cis-male and transgender/non-binary individuals, which will allow examining between-sex/gender differences in JFM. Second, our sample had a low representation of different ethnicities and participants with low socioeconomic status. Community-oriented research is crucial to overcome the predominance of white patients with medium/high socioeconomic status in research samples. Last, this is the first study assessing SFC and the spatial similarity between the SFC pattern and the genetic expression of FM-associated genes in JFM. Thus, our findings should be replicated to determine robustness and translational utility. Importantly, this transcriptomic analysis was exploratory and relied on normative data from the AHBA rather than gene expression measured in our participants. In addition, this resource is based on adult donors (24–57 years), limiting its direct applicability to adolescent populations. Extending this framework to younger populations remains challenging due to the ethical and practical constraints of pediatric tissue collection, as well as the dynamic nature of brain development, which would require spatiotemporal transcriptomic mapping rather than a static atlas.

In conclusion, this study provides the first evidence of the SFC pattern in JFM and the spatial similarity between this pattern and the genetic expression of FM associated genes. Both JFM patients and healthy participants displayed similar SFC patterns. However, JFM patients with higher symptom severity showed reduced connectivity in the second step SFC between bilateral secondary somatosensory cortices and the right anterior insula. Furthermore, there is spatial overlap between our imaging findings and a brain region in the right anterior insula/pars opercularis where a gene previously involved in fibromyalgia, the *MYT1L*, is overexpressed. This study highlights the role of connectomic-genetic analyses in furthering our understanding of the multiple neurophysiological mechanisms contributing to disease symptoms.

## 5. Acknowledgments

This work was supported by Cincinnati Children’s Hospital Medical Center’s Trustee Grant Award (awarded to MLS) and NIH/NIAMS Grants R01 AR074795 (awarded to MLS) and P30 AR076316, the PID2023-147704OB-I00 funded by MICIU/AEI/10.13039/501100011033/FEDER, UE granted to MLS and the CNS2023-14542 funded by MICIU/AEI/10.13039/501100011033 granted to MLS, NextGenerationEU/PRTR granted to MLS and AMIGO Project ‘A multi-ingredient brain function model predicting chronic pain in youth: a window into future well-being’, reference number 101171809, funded by the European Commission through the HORIZON EUROPE programme, European Research Council (ERC) Consolidator Grant granted to MLS. Marina López-Solà, PhD is hired as part of the Serra Hunter Programme of the Generalitat de Catalunya. We would like to express our most sincere gratitude to all participants and their families. The authors have no conflict of interest to declare. The data that support the findings of this study are available from the corresponding author upon reasonable request.

