## Supplementary Material for "Connectomic Flow Across Cortical Sensory Systems in Juvenile Fibromyalgia"

**Supplementary Methods 1: Participants and recruitment Criteria**

**Supplementary Methods 2: Transcriptomic and genetic analyses**

**2.1 Fibromyalgia-associated genes**

**2.2 Allen Human Brain Atlas (AHBA) dataset**

**2.3 Spatial associations between SFC and cortical transcriptomic information**

**Supplementary Methods 3: MRI Acquisition and Preprocessing**

**References**

**Supplementary Methods 1: Participants and recruitment Criteria**

Participants were drawn from the Fibromyalgia Integrative Training program for Teens (FIT-teens) cohort [16]. Juvenile fibromyalgia (JFM) was diagnosed according to established clinical criteria, and healthy controls were recruited from the same community and screened to exclude chronic pain conditions and major medical or psychiatric disorders. Detailed inclusion and exclusion criteria are provided in the original FIT-teens publication.

The sample was restricted to female adolescents due to the higher prevalence of JFM in girls and the composition of the parent clinical trial, in which 86% of eligible participants were cisgender girls. This limited the feasibility of including other genders and helped reduce potential confounding effects related to sex differences in pain processing and brain development [4,18,19].

JFM patients were recruited at the Cincinnati Children's Hospital Medical Center, after being diagnosed by a pediatric rheumatologist or a pain physician using 2010 American College of Rheumatology criteria. Healthy participants were recruited using the Office of Computational and Technical Research (OCTR) participant database of people interested in being notified about studies being conducted at Cincinnati Children's Hospital and from previous studies in which participants consented to be contacted for new research opportunities.

Inclusion criteria were the following: 1) A JFM diagnosis confirmed by a pediatric rheumatologist or pain physician using the 2010 American College of Rheumatology (ACR) criteria adapted for pediatric use. These criteria were chosen because they have been specifically adapted and validated in adolescent females with JFM and were also applied in the parent clinical trial FIT-teens [16]. 2) An average pain intensity of 3 or greater out of 10 over the past week, as measured by the Brief Pain Inventory (BPI) (Daut et al., 1983), and 3) Functional Disability Inventory (FDI) scores higher than 7 out of 60, indicating at least mild disability [17]. Inclusion criteria for healthy controls included 1) being physically and psychologically healthy (i.e., not diagnosed with chronic pain, psychiatric, neurological, or inflammatory disorders), and 2) average pain score of 0 on a Numeric Rating Scale and an FDI score equal or below 7.

Subjects with contraindication to MRI scanning, developmental delay, major neurological or psychiatric disorders (except for JFM patients with mood, anxiety, or obsessive-compulsive disorders undergoing treatment), a positive pregnancy test, or taking opioid medication were not eligible. All participants included in the study were under no medication or under a stable medication regime for a minimum of 3 weeks prior to the MRI assessment.

All participants and their parent/legal guardians provided written informed consent/assent. The Cincinnati Children's Hospital Medical Center's Institutional Review Board approved the study protocol and consent forms (ID: 2017-7771).

**Supplementary Methods 2: Transcriptomic and genetic analyses**

**2.1 Fibromyalgia-associated genes**

A Genome-Wide Linkage Scan (GWLS) study [2] identified one major locus on chromosome 17p11.2–q11.2, which coincides with the chromosomal location of two potential FM candidate genes, *SLC6A4* and *TRPV2*. *SLC6A4* encodes the serotonin transporter (*SERT*); polymorphisms in *SLC6A4* have been previously associated with chronic pain conditions, higher levels of depression, and psychological disorders in chronic pain patients [14,15]. *TRPV2* encodes ion channels activated by high temperatures; its protein is expressed in sensory ganglia and may contribute to altered pain thresholds in FM patients [23].

A Genome-Wide Association Study (GWAS) study [11] found two FM-associated variants, *rs11127292* SNP and an intronic copy number variant link to *MYT1L*, a gene involved in neuronal differentiation by specifically repressing the expression of non-neuronal genes during neuron differentiation and is involved in cognitive disability [25]. Last, *NRXN3* is a cell membrane protein that functions in the nervous system as a receptor involved in cell recognition and adhesion [26].

Together, these genes provide a biologically grounded set of markers for linking fibromyalgia-related genetic variation to spatial patterns of cortical organization derived from the AHBA.

**2.2: Allen Human Brain Atlas (AHBA) dataset**

The Allen Human Brain Atlas (AHBA) [13,22] is a transcriptional atlas of the adult human brain derived from histological analysis and microarray profiling. Data were originally obtained from six donors between 24 and 57 years of age, with no known neuropsychiatric or neuropathological history [13]. The transcriptome dataset consists of genetic expression profiles of 20,787 protein-coding genes extracted from 58,692 measurements from 3,702 brain samples. These samples were initially mapped to native three-dimensional MRI coordinates, then to the Montreal Neurological Institute (MNI) coordinate space. Each donor's closest blood relative provided informed consent for brain tissue collection.

**2.3 Spatial associations between SFC and cortical transcriptomic information**

To investigate cortical gene expression profiles associated with FM, we studied patterns of topographical co-localization between phenotypic FM brain maps (obtained in SFC) and the cortical gene expression values of familiar FM genes (*SLC6A4, TRPV2*, *MYT1L* and *NRXN3*). We leveraged our well-established neurogenetic pipeline, initially developed by Sepulcre [21] and subsequently employed in several neuroscience translational investigations (e.g., Basaia et al., 2022; Bueichekú et al., 2020, 2024; Diez et al., 2024; Diez & Sepulcre, 2021; Gutiérrez-Zúñiga et al., 2022) [5–7,9,10,12]. In our approach, we use the Desikan–Killiany brain ROI atlas to conduct a surface anatomical transformation of the AHBA transcriptome and extract gene expression levels for specific genes and regions. The Desikan–Killiany atlas was selected due to its widely used anatomically defined cortical parcellation, which provides a structurally interpretable and robust framework for mapping gene expression data onto cortical regions. As previously described [1,7,8], to derive the transcriptome data for our custom atlas, we followed three steps for each AHBA individual and obtained a group expression matrix with the median values: (1) expression values from multiple probes were mean averaged for each gene; (2) each sample was mapped to a brain region in which the samples not assigned to any region were evaluated and the samples with a distance <3 mm to any neocortical region was assigned to it; and (3) the median genetic expression across all samples within each cortical region were computed. A group expression map was computed after calculating the median expression values of the six individual donors. Then, using in-house developed MATLAB coding and a region-wise neurogenetic approach, we created single normalized transcriptomic null hypothesis distribution curves (N=20,737 genes) for each of the 68 cortical regions separately, and delimited significantly overexpressed genes per region (threshold > 1.96 SD, corresponding to a p-value < 0.05). Next, for each of our FM candidate genes - *SLC6A4*, *TRPV2, NRXN3* and *MYT1L* – we recorded whether their transcriptomic value fell under the significance area in each cortical region (see **Figure 5**). Importantly, this procedure assesses gene expression relative to a region-specific null distribution rather than performing statistical comparisons across regions. Therefore, no correction for multiple comparisons across cortical regions was applied.

**Supplementary Methods 3: MRI Acquisition and Preprocessing**

T1-weighted gradient-echo sequence images and T2*-weighted gradient echo, echo-planar images (EPI) sequences were collected on 3.0-Tesla MR Philips Ingenia scanner (Philips Healthcare, Best, Netherlands) equipped with a 32-channel head coil at the Cincinnati Children's Hospital Medical Center. The acquisitions covered the whole brain including the entire cerebellum. Brain structural images were acquired using the standard T1-weighted gradient-echo sequence with the following scan parameters: TR = 10 ms, TE = 1.8, 3.8, 5.8, 7.8 ms, field of view = 256x224x200 mm, voxel size = 1x1x1 mm, number of slices = 200, flip angle = 8°, slice orientation = sagittal, and total scan duration = 4.42 min. Resting-state BOLD fMRI data were collected using a T2* weighted echo planar imaging sequence with a multiband sensitivity encoding (SENSE) technique and the following scan parameters: multiband acceleration factor = 4, TR = 650 ms, TE = 30 ms, field of view = 200 mm, flip angle = 53°, voxel size =2.5 x 2.5 x 3.5 mm, slice orientation = transverse (parallel to the orbitofrontal cortex line), slice thickness = 3.5 mm, number of slices = 40 (providing whole-brain coverage), number of volumes = 571, dummy scans = 12, and total scan duration = 6:13 min. Board-certified neuroradiologists reviewed images to ensure participants did not show anatomical abnormalities.

Imaging data were preprocessed using the CONN Toolbox-20.b (www.nitrc.org/projects/conn) running on MatLab version 2021a (Math Works Inc, Natick, MA). The preprocessing followed the standard steps: realignment and unwarping, outlier detection, segmentation and normalization of structural images, and normalization and smoothing of functional images. The first 12 volumes of the BOLD sequence were discarded to allow magnetization to reach equilibrium, and the remaining volumes were realigned to the first one. Based on head-motion parameters estimated during realignment, volumes with framewise displacement >0.9mm or global BOLD signal changes >5 standard deviations were flagged as outliers [20]. Realigned images were co-registered to the T1-weighted volume, which was normalized into standard Montreal Neurological Institute (MNI) space and segmented into gray matter, white matter, and cerebrospinal fluid compartments [3]. The same estimated non-linear transformation was then applied to the functional images. Finally, functional images were smoothed with a 6mm full width at half-maximum Gaussian kernel. In a subsequent temporal denoising step, noise components from white matter and cerebrospinal fluid areas, characterized by the principal component-based aCompCor method, were regressed out from BOLD timeseries along with subject-motion parameters, motion outliers, and a linear detrending term [24]. A band-pass filtering of 0.008 Hz-0.09 Hz was performed. Images were inspected for artifacts at every step and quality assurance plots were visually examined. Global signal change and framewise displacement did not differ between groups ($t = 1.61$, $p = 0.11$; $t = 0.80$, $p = 0.43$, respectively).
